# Molecular Conformer Sampling Beyond Generations

**DOI:** 10.64898/2026.11.19.767840

**Authors:** Alvand Vahediahmar, Aaron Zweig, Mingxuan Zhang, Elham Azizi

## Abstract

Sampling the low-energy conformational states available to a molecule is a fundamental problem in computational chemistry and drug discovery, since molecular properties and downstream design decisions depend on the relevant three-dimensional geometries rather than on a single structure. Recent diffusion and flow-based generative models can efficiently sample conformer candidates from molecular graphs, but under a fixed output budget their samples do not cover the full set of relevant conformational states of a molecule, while practical bench-marks and downstream applications require a small ensemble that is both diverse and physically plausible. We present a post-generation selection framework that turns candidate pools from frozen, pre-trained generators into compact conformer ensembles without any retraining or fine-tuning of the underlying generator. The framework offers two selectors that share the same candidate pool and the same learned reference-likelihood classifier: CovSelect, a coverage-maximizing selector that greedily reaches as many distinct reference states as the pool allows, and RefSelect, a tunable selector that trades some of that coverage for higher precision and, at its balanced setting, matches or exceeds strong generative base-lines on both measures at once. On the full GEOM-DRUGS test set, CovSelect reaches 91.33% mean recall (98.36% median), outperforming every raw generator we compare against, and RefSelect at its balanced setting exceeds the strongest raw baseline, DMT-L, on both recall and precision. Finally, across several out of distribution benchmarks (QMugs, pepconf, and Platinum), CovSelect substantially outperforms all raw generative baselines in recall and precision.

## 1 Introduction

Modern generators can produce far more conformers than a downstream workflow can inspect. If we oversample generously, for example by sampling 16 times more candidates than the number of ground truth conformers and pooling them on the GEOM-DRUGS test set, the resulting candidate pool contains virtually every conformer in the whole test set and would count as a complete hit under standard benchmarks, with mean recall of 97% and median recall of 100%. This retrospective *pool ceiling* is measured with reference coordinates after generation. It shows that candidate generation and fixed-budget ensemble construction are distinct problems. We therefore introduce a framework that separates these stages: oversample from frozen generators, infer where useful reference states are likely to lie, and then construct a compact ensemble of exactly *B* = 2*n* conformers, where *n* is the reference count recorded in its input file (Axelrod & Gómez-Bombarelli, 2022). Passing the full pool to an evaluator would violate that protocol and multiply the cost of the downstream scoring, refinement, or experimental checks that make up the truly costly part of the whole pipeline, rather than the wall-clock time spent on a GPU. We ask how to choose the most useful *B* conformers from a pool produced by frozen generators.

The limited budget makes coverage especially important. If no returned conformer represents a particular state, later checks cannot recover it. Recovering as many states as possible is therefore more valuable than returning samples that already sit exactly on a reference structure, since a downstream relaxation step can reach a nearby reference from a close generated sample. The intuition behind this work is that a large generated pool of candidates can cover almost all such states, but current models are evaluated under a strict per-molecule budget, and when they sample only *B* candidates directly they miss many valid states. To address this, we separate generation from selection and follow an *oversample first, then select* pipeline: build a large candidate pool once, then choose the *B* conformers to submit. A selection rule must therefore balance how much each candidate adds to the ensemble against evidence that it resembles a reliable conformer, evidence we learn once with a classifier (Section 4) and never retrain at test time. Figure 1 shows a held-out example in a fixed, train-fitted conformer embedding. This paper makes three contributions:

**Figure 1:**
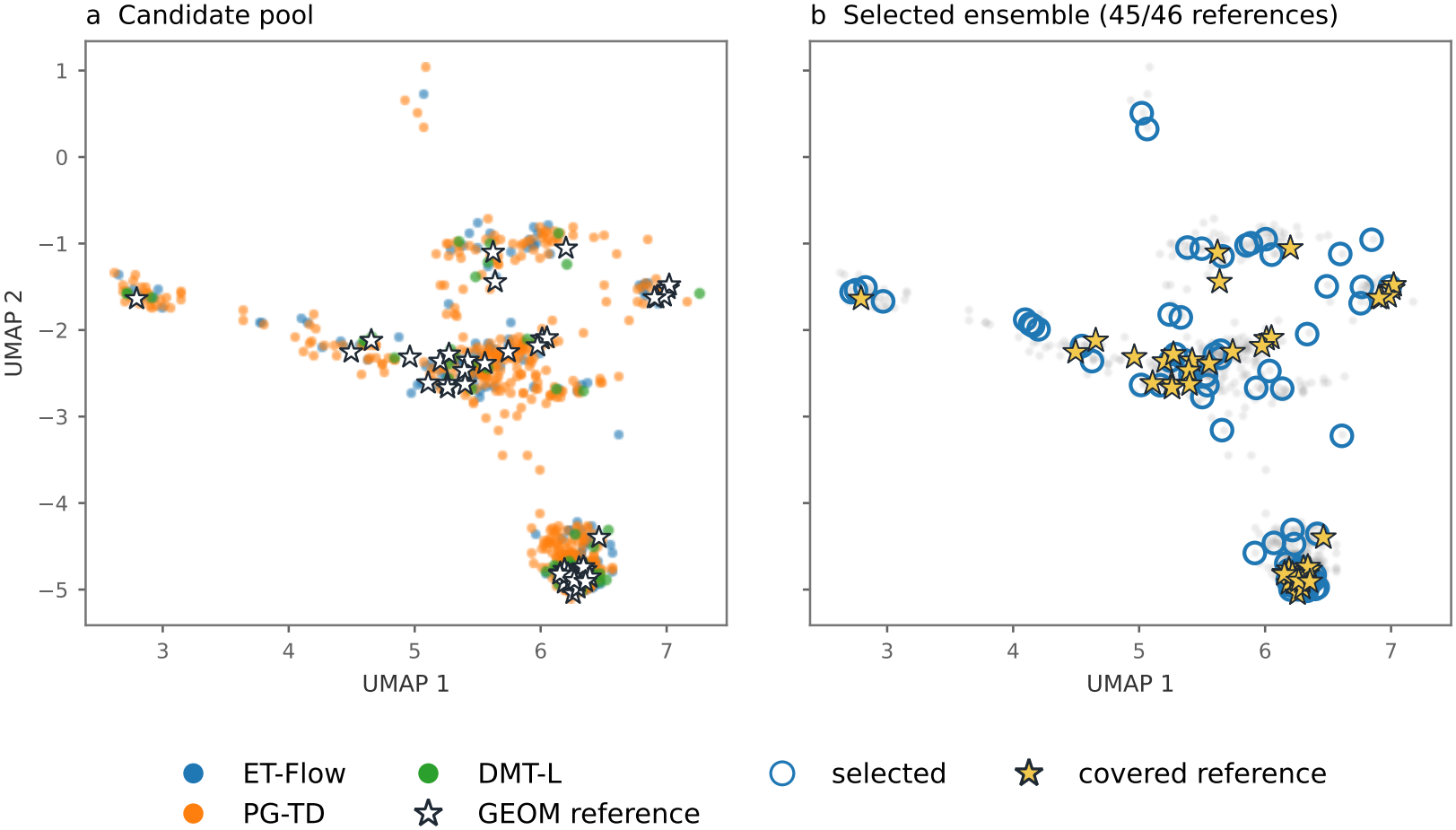
Held-out molecule U7 (SMILES `CC(C)(C)CC(C)(C)NC(=O)Nc1cccc([N+](=O)[O-])c1`) projected into a UMAP fitted on 2,000 disjoint training molecules. The plot shows 736 pool candidates, 92 selected conformers, 46 references, and 92 separate DMT-L samples. Rings mark selected candidates and stars mark references reached by exact RMSD, 45 of 46 at 0.75 Å .

- **A framework for fixed-budget ensemble construction**. We separate conformer generation from ensemble selection, constructing compact sets of exactly *B* = 2*n* conformers from oversampled outputs of frozen generators without retraining them.
- **Latent reference-field inference**. We combine learned candidate reference-match probabilities, a training-derived coverage kernel, and nonnegative deconvolution to estimate reference mass over candidate locations, using the benchmark-provided reference count but no test reference coordinates.
- **Redundancy-aware selection with tunable coverage and precision**. Greedy selection combines estimated reference coverage, a precision proxy, and a training-estimated redundancy correction. CovSelect targets coverage, while RefSelect balances coverage and precision through a single weight. On GEOM-DRUGS, CovSelect achieves 98.36% median recall, and RefSelect at *µ* = 0.5 exceeds next best baseline on both recall and precision.

## 2 Related Work

### Conformer generation

GEOM provides the energy-annotated conformer ensembles used in many generative benchmarks (Axelrod & Gómez-Bombarelli, 2022). Torsional diffusion models flexibility in the molecule’s torsion angles (TD) (Jing et al., 2022), and particle guidance changes how its samples interact during generation to improve ensemble diversity (PG TD) (Corso et al., 2023). ET-Flow instead uses equivariant flow matching (Hassan et al., 2024), and NExT-Mol’s diffusion molecular transformer supplies the DMT-L baseline (Liu et al., 2025). These methods principally improve the proposal distribution. We use their frozen outputs as candidate conformers and study which candidates should be returned under a common budget.

### Conformer-set reduction

Selecting representative conformers after generation has a longer history. PubChem3D clusters a large generated set by RMSD before retaining a manageable ensemble, and this reduction can lose proximity to experimentally observed ligand shapes (Kim et al., 2013). Automated RMSD-based clustering likewise compresses candidate ensembles for drug modeling (Kim et al., 2017). Classical facility-location selection formalizes coverage of observed candidates (Nemhauser et al., 1978). Our objective instead estimates coverage of reference states from candidate evidence learned on other molecules, then balances it with a candidate-level precision proxy. Its inverse-field calculation is adapted from Richardson–Lucy deconvolution (Richardson, 1972; Lucy, 1974).

## 3 Task and Evaluation

For each molecule, let *n* be the reference count recorded in the benchmark input file, and let *G* = {*g*_1_, …, *g*_*n*_} be the reference conformers retained by the evaluator’s cleaning step. The selector receives *B* = 2*n* as its return budget, and it never sees *G*. Let *C* = {*c*_1_, …, *c*_*N*_ } be candidates, *d* symmetry-corrected heavy-atom RMSD, and *τ* = 0.75 Å . A selected set *S* ⊆ *C* has |*S*| = *B*. Recall is |*G*|^−1^ ∑_*g*∈*G*_ **1**[min_*s*∈*S*_ *d*(*g, s*) *< τ* ], and precision is *B*^−1^ ∑_*s*∈*S*_ **1**[min_*g*∈*G*_ *d*(*g, s*) *< τ* ]. Recall and precision average minimum RMSD (AMR) replace each indicator with its corresponding minimum RMSD, and lower AMR is better. We aggregate each metric over molecules and report its mean and median.

## 4 Methodology

Our selection pipeline has three stages, shared by both selectors. We first build a large pool of candidate conformers per molecule from frozen, pre-trained generators (Section 4.1). We then score each candidate with a classifier, trained once on separate molecules and never fine-tuned, and use that score as a per-candidate quality signal, a tie-breaker among otherwise similar candidates rather than a hard filter (Section 4.2). Finally, we combine the classifier’s scores with a coverage kernel and a reference field, an estimate of where the unseen reference conformers most likely sit given the candidate pool, to greedily select exactly *B* candidates under a fixed-budget objective (Sections 4.3 and 4.5). The redundancy correction in Section 4.4 refines this objective so that near-duplicate candidates are not double-counted toward coverage.

### 4.1 Candidate Pool

The full-cohort experiment reuses an archived multi-source candidate pool. A raw-coordinate audit of its 999 molecules, performed after finite-coordinate filtering (discarding any candidate whose atomic coordinates are not finite numbers, which happens when a generator’s sampler fails on a difficult molecule), verifies two contiguous source segments: nominally 8 × ET-Flow and 8 × PG-TD. One molecule has unequal source counts, so the multipliers describe the recipe rather than an exact per-molecule count. DMT-L is evaluated as an external raw-generator baseline, and it is not in this pool. The selector filters non-finite coordinates, computes candidate-candidate RMSD, and returns distinct candidate indices. The returned sets were audited for duplicate coordinates. This pool is a collection of alternative coordinate sets for the *same* molecular graph, and it is never itself the reported answer. It gives the selector options while the final *B* positions remain fixed. Inference features are computed before looking at any test reference.

### 4.2 Reference Likelihood Model

Selection needs a way to tell, for each candidate, how likely it is to actually match one of the (hidden) reference conformers. We learn that likelihood once, on training molecules, with a frozen classifier. On GEOM training molecules, each generated candidate receives a label *y*_*i*_ = **1**[min_*g*∈*G*_ *d*(*c*_*i*_, *g*) *< τ* ], meaning it is within *τ* of some real reference conformer for that molecule. A histogram gradient-boosted tree is fit to predict *p*_*i*_ = Pr(*y*_*i*_ = 1 | *z*_*i*_), the probability of that label, from features *z*_*i*_ available at proposal time, before any reference is used: how many other candidates for the same molecule sit within several RMSD radii of candidate *i* (a local density feature, capturing whether *i* sits in a crowded or a sparse region of the pool), where *i* ranks among the pool by predicted energy, and summaries of the predicted energy and force on *i*’s immediate atomic neighborhood. The energy and force values behind these features come from two frozen, off-the-shelf machine-learned interatomic potentials, AceFF (Farr et al., 2026) and AIMNet2 (Anstine et al., 2024), applied to each candidate’s raw coordinates. Molecule size is also included as a feature. The model is fitted on 164,312 proposals from 2,000 unique training molecules, with five molecule-grouped validation folds, so every proposal for one training molecule stays in a single fold and no test connectivity identity overlaps a training one. The learned signal is a probability of *matching a released GEOM conformer*, not a complete physical-validity judgment, and Appendix A gives the full feature list and training details.

### 4.3 Coverage Kernel and Reference Field

The classifier gives a per-candidate score, but selection needs to know how many distinct reference states the pool actually contains and where they are likely to be, so that the selector can spread its picks across them instead of clustering on one lucky region. We address this with two calibrated, train-only quantities: a coverage kernel that converts geometric distance into a probability of coverage, and a reference field that redistributes the classifier’s scores into an estimate of where the hidden references sit.

On training molecules we estimate *κ*(*u*), the chance that a submission covers a reference conditional on that reference being within *τ* of its nearest pool candidate, when the submission is distance *u* from that nearest candidate. This conditioning separates geometric displacement from whether the pool reaches a reference at all. The kernel decreases with distance and has finite measured support. We place nonnegative latent reference mass *λ*_*j*_ at candidate *c*_*j*_, representing our estimate of how much of the (unseen) reference set sits near that candidate. Under a Poisson approximation, the probability of a candidate being near at least one reference is

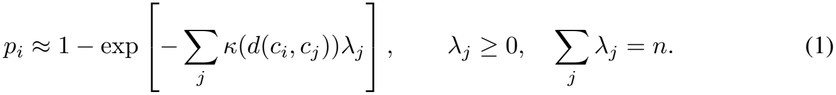

Here *p*_*i*_ is the classifier’s direct estimate for candidate *i*, based only on its available features. We treat these scores as blurred observations of an underlying reference field and use −log(1 − *p*_*i*_) followed by thirty nonnegative Richardson-Lucy updates to infer *λ*, while enforcing total mass *n* = *B/*2. We then compute a probability from this inferred field,

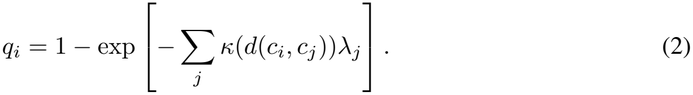

Unlike *p*_*i*_, which is a local classifier score, *q*_*i*_ is a field-based coverage probability that uses the inferred reference mass across the entire pool through *K*_*ij*_. Thus, *p*_*i*_ provides the local evidence for inferring the reference field, while *q*_*i*_ measures how well candidate *i* covers that field. The selector uses *q*_*i*_, rather than *p*_*i*_, in its quality term.

### 4.4 Redundancy Calibration

The reference field tells the selector where references likely are, but it does not yet know that two nearly identical candidates usually stand or fall together: if one matches a reference, the other almost certainly does too, so returning both wastes budget that could have covered a different state. We correct for this with a redundancy calibration.

Call a returned candidate a *witness* if it is close enough to a reference to plausibly count toward that reference’s coverage. The field-smoothed *q*_*i*_ estimates whether returning candidate *i* would count as a witness toward precision, with the learned *p*_*i*_ as its input through Eq. 1. For recall, two nearly identical witnesses usually succeed or fail together, since a small shift that misplaces one is likely to misplace an almost identical neighbor the same way, while a witness on another side of the same possible reference can still rescue it. On disjoint training molecules we estimate *ρ*(*v*), the conditional extra chance of covering a reference that a second witness provides, given that it is separated by distance *v* from the nearest already-chosen witness. Its measured value is about 0.20 below 0.25 Å and 0.83 between 0.25 and 0.50 Å, and estimates above one for larger separations are clipped to independence, since a distant second witness cannot be more than fully informative. This lets the selector discount near-duplicate picks while still crediting distinct nearby candidates that genuinely improve coverage.

### 4.5 Fixed-Budget Selection

The previous three components, the classifier’s score, the reference field, and the redundancy correction, combine into a single objective that the selector optimizes greedily under the fixed budget *B*. Without the redundancy correction, the estimated objective is

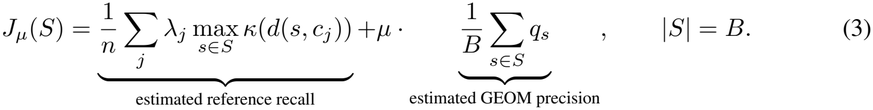

Larger *µ* favors higher predicted precision over additional coverage, and this single weight is what lets RefSelect move along the recall-precision trade-off. CovSelect corresponds to the coverage-only endpoint of this same objective. Greedy selection adds the candidate with the largest marginal objective increase until exactly *B* are returned, one candidate at a time, so the same candidate can become redundant once a nearby one is selected, without any hand-written rule reserving a fixed number of slots for diversity.

The deployed objective replaces the first term with a correlated-witness update using *ρ* from Section 4.4. The standard (1 − 1*/e*) guarantee applies to the nonnegative facility-location form in Eq. 3, though no such guarantee has been proved for the deployed nearest-witness correction. Selection uses only the candidate pool, the frozen models, and the permitted count *B*. The complete inference procedure is summarized in algorithm 1. It applies the classifier, deconvolves its scores into the reference field, and then greedily selects exactly *B* candidates using the coverage-quality objective with the redundancy correction.

## 5 Experimental Setup

### Dataset and metrics

The common evaluable GEOM-DRUGS test set has 999 molecules. Candidate labels, kernel calibration, and redundancy calibration use molecule-disjoint GEOM training data. The kernel study uses 2,000 training molecules and 36,306 conformers. We use 20,000-draw molecule-paired bootstrap intervals for method differences, resampling molecules rather than individual conformers. We reproduce raw ET-Flow, PG-TD, and DMT-L ensembles at the 2 × return count. CovSelect, RefSelect, and uniform random select from a 16 × pool. The random row averages ten fixed-seed draws per molecule.

## 6 Results

### 6.1 GEOM-DRUGS Performance

Table 1 and Figure 2 show how the returned ensemble changes as RefSelect increases the weight on reference likelihood. CovSelect achieves the highest recall in the table, with 91.33% mean and 98.36% median, while RefSelect provides a tunable recall-precision frontier. At *µ* = 0.5, it exceeds DMT-L by 2.83 recall points and 3.33 precision points, while at *µ* = 1.2 it exceeds ET-Flow by 3.81 recall points and 1.67 precision points. CovSelect’s 91.33% recall comes at lower preci-sion (55.52%) because its coverage objective spreads selections across the surplus pool, recovering reference states missed by raw outputs while also selecting more non-matching candidates. Ref-Select at *µ* = .2 achieves nearly the same recall with higher precision. As a same-pool control, uniform selection gives 80.17% recall and 66.20% precision, while ranking only by the frozen likelihood gives 44.09% recall and 83.66% precision, showing the cost of concentrating selections in a few regions. These results show that the gains come from selection rather than simply increasing the candidate pool, motivating the question of whether unguided geometric selection on the same pool can achieve comparable performance.

**Table 1:**
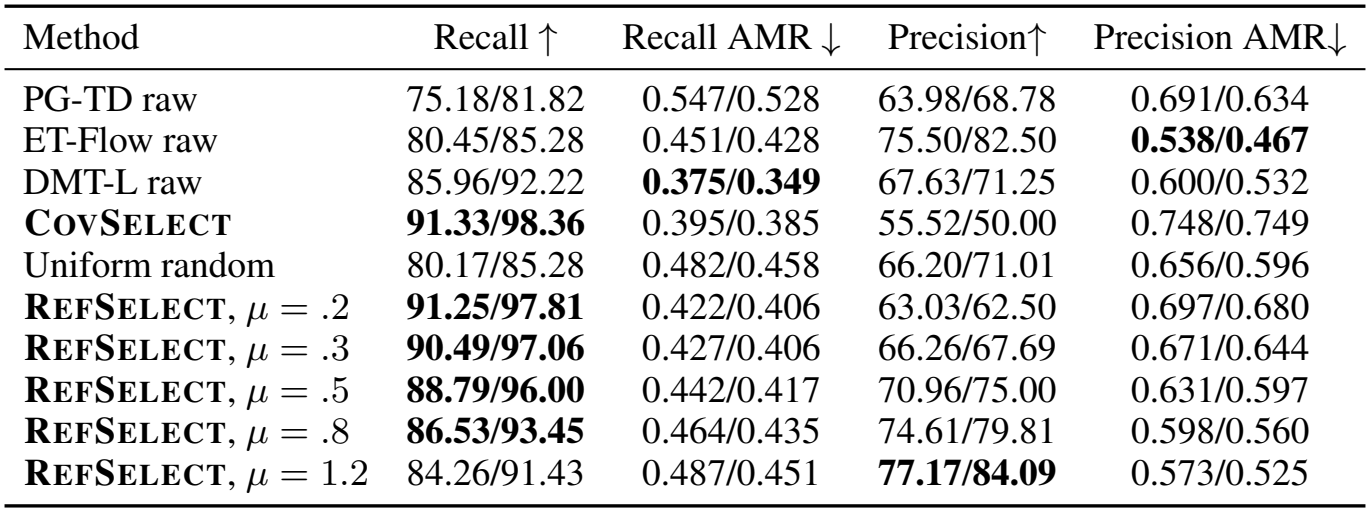
GEOM-DRUGS recall and precision at the 0.75 Å threshold, and their average minimum RMSD (AMR, Å, lower is better), on the full test set. Each cell is mean/median across molecules. Every method returns *B* = 2*n* conformers. PG-TD’s AMR columns use one fewer molecule (one nonfinite result). Bold marks the best mean and the best median in each column.

| Method | Recall ↑ | Recall AMR ↓ | Precision↑ | Precision AMR↓ |
|---|---|---|---|---|
| PG-TD raw | 75.18/81.82 | 0.547/0.528 | 63.98/68.78 | 0.691/0.634 |
| ET-Flow raw | 80.45/85.28 | 0.451/0.428 | 75.50/82.50 | **0.538/0.467** |
| DMT-L raw | 85.96/92.22 | **0.375/0.349** | 67.63/71.25 | 0.600/0.532 |
| **COVSELECT** | **91.33/98.36** | 0.395/0.385 | 55.52/50.00 | 0.748/0.749 |
| Uniform random | 80.17/85.28 | 0.482/0.458 | 66.20/71.01 | 0.656/0.596 |
| **REFSELECT**, μ = .2 | **91.25/97.81** | 0.422/0.406 | 63.03/62.50 | 0.697/0.680 |
| **REFSELECT**, μ = .3 | **90.49/97.06** | 0.427/0.406 | 66.26/67.69 | 0.671/0.644 |
| **REFSELECT**, μ = .5 | **88.79/96.00** | 0.442/0.417 | 70.96/75.00 | 0.631/0.597 |
| **REFSELECT**, μ = .8 | **86.53/93.45** | 0.464/0.435 | 74.61/79.81 | 0.598/0.560 |
| **REFSELECT**, μ = 1.2 | 84.26/91.43 | 0.487/0.451 | **77.17/84.09** | 0.573/0.525 |

**Figure 2:**
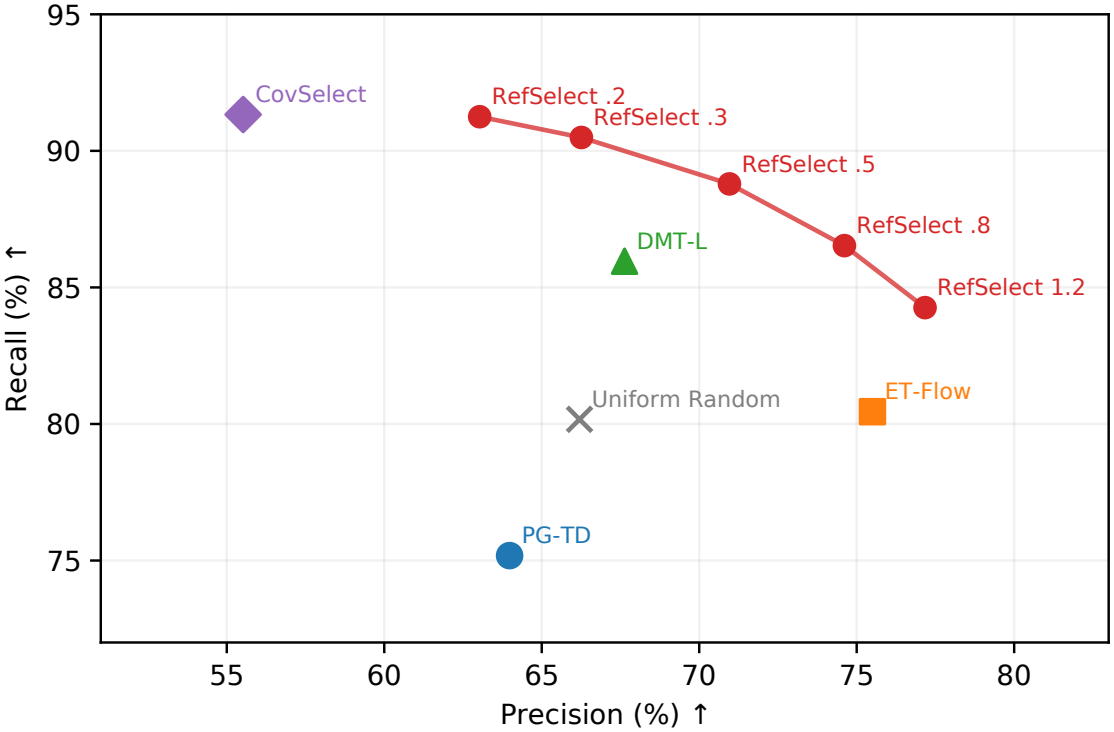
Recall and precision on GEOM-DRUGS at the same returned count. Random selection averages ten draws per molecule

#### Algorithm 1

RefSelect: learned fixed-budget conformer selection

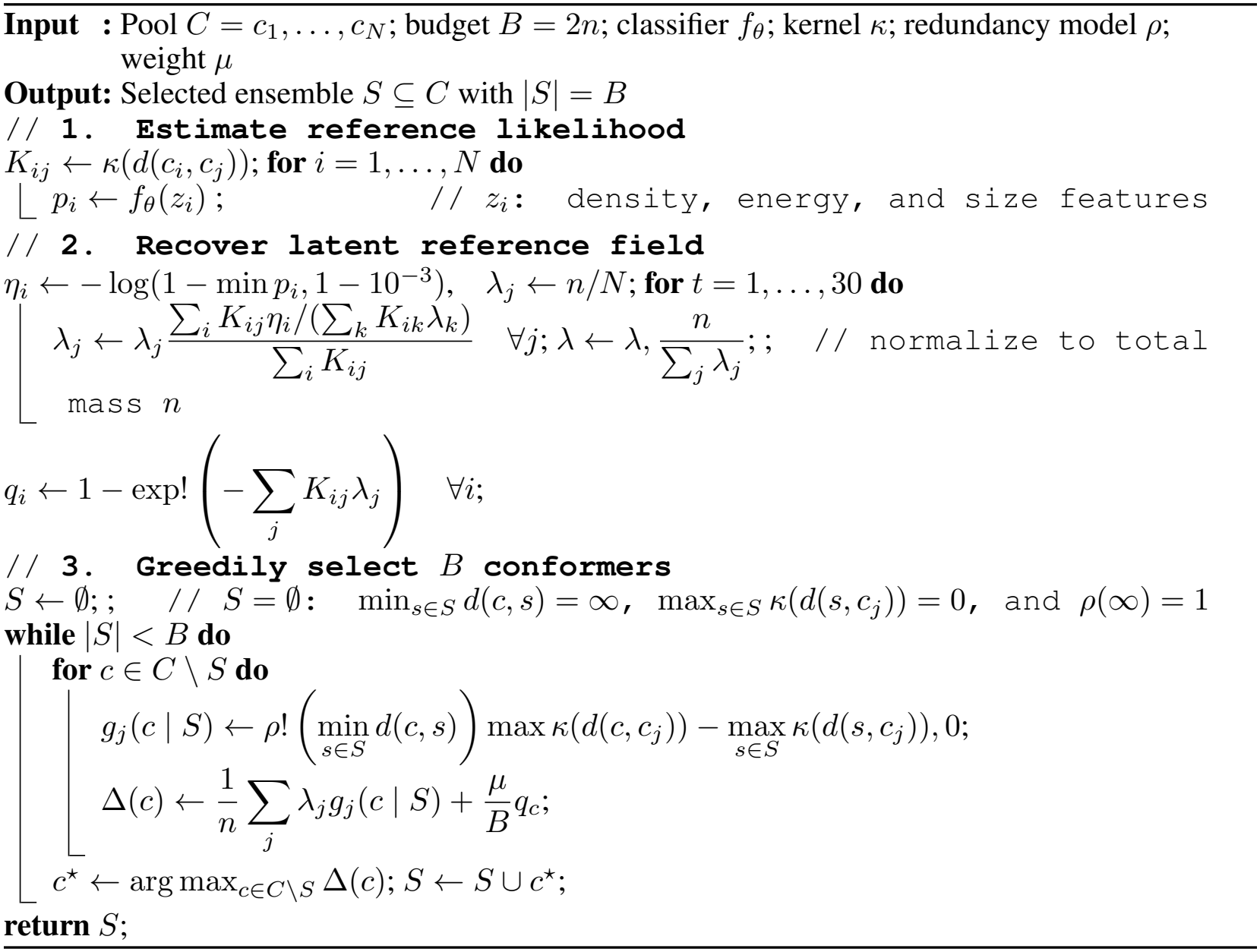

### 6.2 Classical Diversity Selection

To evaluate whether unguided geometric heuristics suffice, we compare our framework against two classical diversity rules on identical pool at budget *B* = 2*n*: *Farthest-Point Sampling* (FPS) (Gon-zalez, 1985) and *k-medoids clustering* (*k* = *B*) (Kaufman & Rousseeuw, 1990). Both rules operate strictly on the symmetry-corrected heavy-atom RMSD matrix without access to likelihood scores, kernels, or reference field priors. As shown in Table 2, *k*-medoids achieves strong Recall AMR (0.405 Å), but falls well short of CovSelect’s peak recall (91.33%) and RefSelect’s balanced precision-recall trade-off. FPS over-indexes on peripheral outliers, severely hurting precision (42.18%). These findings demonstrate that learned reference-likelihood and deconvolved field estimates are essential for isolating physically plausible states beyond simple spatial dispersion. Having established the necessity of learned selection on GEOM-DRUGS, we next evaluate how effectively these models generalize to out-of-distribution chemical spaces.

**Table 2:** Selection rules on the identical same candidate pool at *B* = 2*n* (mean/median across molecules). Uniform random averages ten draws per molecule.

| Method | Recall ↑ | Recall AMR ↓ | Precision ↑ | Precision AMR ↓ |
|---|---|---|---|---|
| Farthest-point | 80.34 / 97.07 | 0.512 / 0.495 | 42.18 / 35.71 | 0.918 / 0.881 |
| k-medoids | 86.56 / 93.00 | **0.405 / 0.380** | 66.76 / 71.43 | 0.646 / 0.593 |
| Uniform random | 80.17 / 85.44 | — | 66.20 / 71.42 | — |
| **REFSELECT** (*μ*=0.3) | **90.66 / 97.15** | 0.425 / 0.406 | 66.52 / 67.86 | 0.669 / 0.641 |
| **REFSELECT** (*μ*=0.5) | **88.96 / 96.06** | 0.440 / 0.413 | **71.21 / 75.00** | **0.629 / 0.595** |

### 6.3 Out-of-distribution Evaluation

Every model in this section is used exactly as released, with no training or fine-tuning of any kind, including for CovSelect: unlike its row in Table 1, the CovSelect used here has no energy or quality tie-break and no prior calibrated on GEOM training molecules, so its advantage below comes entirely from the selection rule, not from anything fit on drug-like chemistry. We first ran the frozen generators on three chemically different test sets, pepconf(Prasad et al., 2019), QMugs (Isert et al., 2022), and Platinum (Friedrich et al., 2017), none of which overlap GEOM-DRUGS, under the same 2*n* return-count rule used everywhere else. Table 3 reports recall and precision (Appendix B reports the same comparison in AMR). CovSelect wins recall by a wide margin over the raw generators on every set, and RefSelect pushes precision further still, winning it outright on pepconf and Platinum and matching CovSelect’s recall lead on Platinum too, with DMT-L the weakest model throughout, consistent with a generator tuned on drug-like GEOM-DRUGS chemistry and then applied zero-shot with no OOD-specific selection step at all.

**Table 3:**
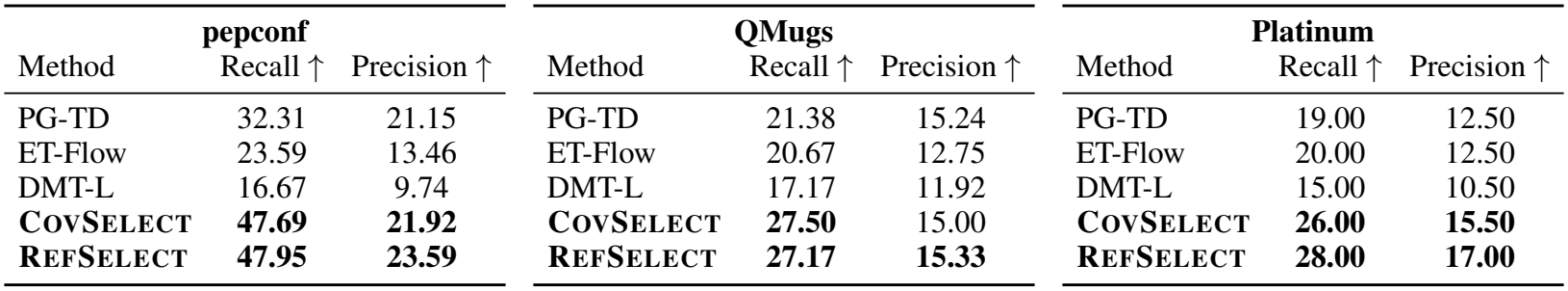
Out-of-distribution recall and precision (%) at the 0.75 Å threshold, at the matched 2*n* returned count, mean across each set’s molecules (pepconf, QMugs, and Platinum do not overlap GEOM-DRUGS). PG-TD and DMT-L are raw generators, CovSelect selects from the union of the PG-TD and ET-Flow pools with no GEOM-trained prior.

| **pepconf** |  |  | **QMugs** |  |  | **Platinum** |  |  |
|---|---|---|---|---|---|---|---|---|
| Method | Recall ↑ | Precision ↑ | Method | Recall ↑ | Precision ↑ | Method | Recall ↑ | Precision ↑ |
| PG-TD | 32.31 | 21.15 | PG-TD | 21.38 | 15.24 | PG-TD | 19.00 | 12.50 |
| ET-Flow | 23.59 | 13.46 | ET-Flow | 20.67 | 12.75 | ET-Flow | 20.00 | 12.50 |
| DMT-L | 16.67 | 9.74 | DMT-L | 17.17 | 11.92 | DMT-L | 15.00 | 10.50 |
| **CovSelect** | **47.69** | **21.92** | **CovSelect** | **27.50** | 15.00 | **CovSelect** | **26.00** | **15.50** |
| **RefSelect** | **47.95** | **23.59** | **RefSelect** | **27.17** | **15.33** | **RefSelect** | **28.00** | **17.00** |

#### Independently generated references

pepconf, QMugs, and Platinum still reuse conformer references computed by the same kind of pipeline used to build GEOM-DRUGS, so their references and our test molecules share a common origin despite being chemically different. To test our selectors against references with no shared origin, we picked four molecules whose scaffolds do not occur in GEOM-DRUGS, and generated their reference conformers from scratch with CREST (Pracht et al., 2024). Reference count varies from 2 to 774 for highly flexible molecule, which will serve as stress test for the fixed 2*n* budget rule. Table 4 reports each model’s recall and precision on each molecule. CovSelect has the best or tied-best recall on all four molecules, including 100% on three, and the best mean recall at 96.88%, along with best precision on the most flexible molecule. Together with the pepconf, QMugs, and Platinum results, these results show that our selection framework’s advantage is not an artifact of GEOM-DRUGS-like chemistry, GEOM-style references, or classifier training on GEOM-DRUGS, and persists against conformers generated independently on unseen scaffolds.

**Table 4:** Recall/precision (%) at the matched 2*n* returned count on four molecules with independently generated CREST references, none of which occur in GEOM-DRUGS. Bold marks the best value in each row.

| Molecule | PG-TD | ET-Flow | DMT-L | **CovSelect** |
|---|---|---|---|---|
| Bortezomib (*n*=320) | 78.12 / 57.97 | 51.56 / 49.84 | 81.25 / **33.75** | **87.50** / 27.03 |
| Tavaborole (*n*=2) | 100 / 100 | 100 / 100 | 100 / 100 | **100 / 100** |
| Flusilazole (*n*=16) | 87.50 / 81.25 | 87.50 / 75.00 | 87.50 / **93.75** | **100** / 75.00 |
| 18-Crown-6 (*n*=774) | 74.55 / 97.87 | 99.48 / 99.42 | **100** / 99.68 | **100 / 99.81** |
| Mean | 85.04 / 84.27 | 84.64 / 81.07 | 92.19 / 81.79 | **96.88** / 75.46 |

### 6.4 Return-budget Sensitivity

The benchmark budget is tied to the recorded reference count *n*, and reducing it to *n* (1×) or 0.5*n* (0.5 ×) changes the decision problem without retraining any model. Submitting fewer conformers is also a stricter evaluation, since there is less room to pad toward a lucky match. We evaluate every method at all three budgets on the identical test set and candidate pools. Table 5 reports coverage and AMR, and Appendix C reports the RefSelect *µ*-grid at 1 × and 0.5× (Table 7). DMT-L is the next-best model at every budget, while CovSelect’s recall advantage grows sharply as the budget tightens: +5.37 points at 2× to +14.75 at 0.5 ×. Its precision deficit also shrinks, from -12.11 points at 2 × to just -0.12 at 0.5 ×, essentially closing the gap while retaining nearly a 15-point recall lead. CovSelect is also the only method whose precision rises as the budget tightens, while raw generators remain essentially flat (Table 5). This follows directly from its greedy coverage-plus-quality objective (Eq. 3): early picks provide the largest marginal coverage at the highest estimated quality, while later picks fill larger budgets by entering progressively less certain regions. A smaller budget therefore retains this confident prefix and raises precision through the selection order itself, without changing the pool or model. RefSelect shows the same pattern across its *µ* grid at both budgets (Table 7): every setting’s precision rises and recall falls from 1 × to 0.5× ; for example, *µ* = 1.2 moves from 76.31/78.57 to 65.59/79.95.

**Table 5:** Recall and precision at 0.75 Å, and their AMR (Å, lower is better), across return budgets on the full test set. Each cell is mean/median. 2× rows repeat Table 1, 1×/0.5× raw rows subsample the same 2× pool, and CovSelect/RefSelect re-select at the smaller count.

| Method | Budget | Recall↑ | Recall AMR↓ | Precision↑ | Precision AMR↓ |
|---|---|---|---|---|---|
| PG-TD raw | 2× | 75.18/81.82 | 0.547/0.528 | 63.98/68.78 | 0.691/0.634 |
| ET-Flow raw | 2× | 80.45/85.28 | 0.451/0.428 | 75.50/82.50 | **0.538**/0.467 |
| DMT-L raw | 2× | 85.96/92.22 | **0.375/0.349** | 67.63/71.25 | 0.600/0.532 |
| **CovSelect** | 2× | **91.33/98.36** | 0.395/0.385 | 55.52/50.00 | 0.748/0.749 |
| **RefSelect**, μ=.5 | 2× | **88.79/96.00** | 0.442/0.417 | 70.96/75.00 | 0.631/0.597 |
| PG-TD raw | 1× | 64.27/66.67 | 0.659/0.634 | 63.77/68.13 | 0.692/0.639 |
| ET-Flow raw | 1× | 69.68/71.74 | 0.583/0.553 | 75.16/83.09 | 0.540/**0.462** |
| DMT-L raw | 1× | 75.38/80.56 | 0.516/0.484 | 67.56/71.74 | 0.601/0.532 |
| **CovSelect** | 1× | **85.73/93.44** | 0.505/0.493 | 58.78/56.41 | 0.730/0.721 |
| **RefSelect**, μ=.5 | 1× | **82.53/89.66** | 0.531/0.496 | 72.72/75.86 | 0.620/0.578 |
| PG-TD raw | 0.5× | 50.60/50.00 | 0.801/0.778 | 63.55/68.75 | 0.698/0.644 |
| ET-Flow raw | 0.5× | 56.90/55.77 | 0.738/0.710 | 75.13/**83.33** | 0.543/0.472 |
| DMT-L raw | 0.5× | 60.72/61.70 | 0.683/0.642 | 68.08/73.33 | 0.597/0.526 |
| **CovSelect** | 0.5× | **75.47/80.93** | 0.632/0.591 | 67.96/68.57 | 0.655/0.631 |
| **RefSelect**, μ=.5 | 0.5× | **71.18/75.00** | 0.661/0.613 | **75.70**/79.55 | 0.594/0.556 |

## 7 Limitations

Our comparisons hold the returned count fixed, but the proposal budgets behind that count differ substantially across methods, and this is the main caveat to keep in mind when reading our results. Raw baselines generate only 2 × conformers, while CovSelect and RefSelect draw a large pool, each followed by feature scoring, pairwise RMSD calculation, and greedy selection, so the larger pools require materially more generation and selection compute than the raw baselines do. A second limitation is the reference-likelihood classifier itself. It is a single histogram gradient-boosted model trained on one fixed feature set, and while it is frozen and never fine-tuned on test data, we have not explored richer geometric descriptors, learned embeddings, or larger training pools that could plausibly sharpen its estimates and, in turn, raise both selectors’ ceiling further. These results indicate that a substantial part of the gap between an oversampled candidate pool and a compact returned ensemble can arise from ensemble construction rather than proposal generation alone.

## 8 Conclusion

This paper argues that selecting a compact conformer ensemble deserves as much attention as generating the candidates it is drawn from. The archived candidate pool reaches 97.25% mean and 100% median reference recall when viewed in full, yet the evaluator only accepts 2*n* conformers, and closing that gap between what a pool contains and what a small ensemble recovers is exactly the problem our two selectors address. CovSelect shows how far a purely coverage-maximizing choice can go at that return count, reaching 91.33% mean recall and a 98.36% median, ahead of every raw generator we tested, on GEOM-DRUGS as well as on OOD samples. RefSelect shows that this coverage advantage does not have to come at precision’s expense: at *µ* = 0.5 it beats the strongest raw generator on both recall and precision simultaneously, and its tunable objective exposes a practical recall-precision trade-off instead of forcing one fixed operating point on every downstream use. Together, these results support fixed-return selection as a genuinely effective way to get more value out of generated candidates, without reinventing the wheel. The larger candidate pools and pairwise selection steps do add compute, and some of our operating points were explored on the reported cohort, so measurements at matched generation cost, on further independent test cohorts, and with direct physical and thermodynamic checks would help determine exactly where this method is most useful in deployment. We see the present results, spanning a standard benchmark, three additional chemically distinct test sets, and four molecules verified against an entirely independent conformer search method, as strong evidence that the selection stage is where a meaningful part of the remaining benchmark gap actually lives.

### Reproducibility statement

The code implements train-only labels, calibration, reference-field inference, exact-budget selection, and paired evaluation. The archived pool has a coordinate-verified source manifest, and a final release should additionally include generator invocation and configuration, checkpoint manifests, and wall-time accounting.

## AI Use Statement

AI tools assisted with drafting and figure design. The authors are responsible for checking all mathematical statements, citations, experiment records, and final text.

## A Reference Likelihood Model Details

This appendix expands Section 4.2 with the full feature list and training protocol behind the frozen reference-likelihood classifier used by both selectors.

### Features

For each candidate conformer *c*_*i*_, proposed for molecule *m* by one of the frozen generators, the classifier sees only quantities computable before any reference conformer for *m* is ever examined.

- **Local candidate density**. For several RMSD radii, the number of other candidates for the same molecule that fall within that radius of *c*_*i*_. A candidate in a crowded region of the pool, where many other candidates propose a similar shape, is treated differently from an isolated candidate, since crowding is itself evidence about how often a generator returns to that region.
- **Within-pool energy rank**. Where *c*_*i*_ ranks, by predicted potential energy, among all other candidates proposed for the same molecule. Lower-energy candidates are, all else equal, more likely to correspond to a real, physically populated conformer.
- **Neighborhood energy and force summaries**. Summary statistics (such as the mean and spread) of the predicted energy and of the predicted atomic forces in *c*_*i*_’s immediate local geometry. Large predicted forces indicate a geometry that is far from a local energy minimum, and are informative independently of the raw energy rank above.
- **Molecule size**. The heavy-atom count of the molecule, since the density and rank features above are naturally read differently for a small, rigid molecule than for a large, flexible one.

The energy and force values behind the second and third feature groups are computed by applying two frozen, off-the-shelf machine-learned interatomic potentials, AceFF and AIMNet2 (Anstine et al., 2024), directly to each candidate’s raw three-dimensional coordinates. Neither potential is fine-tuned on GEOM data, and neither ever sees a reference conformer.

### Labels and training protocol

On GEOM training molecules only, each generated candidate *c*_*i*_ receives a binary label *y*_*i*_ = **1**[min_*g*∈*G*_ *d*(*c*_*i*_, *g*) *< τ* ], indicating whether it lies within the coverage threshold *τ* of some real reference conformer for that molecule. A histogram gradient-boosted tree is fit to predict Pr(*y*_*i*_ = 1|*z*_*i*_) from the feature vector *z*_*i*_ above. Fitting uses 164,312 proposals from 2,000 unique training molecules, split into five molecule-grouped validation folds, so that every proposal belonging to a given training molecule stays in a single fold and is never split across the train and validation sides of that fold. This is the same reason connectivity identities never overlap between our training molecules and any test molecule reported in this paper: the model must generalize to molecular graphs, not just to conformers of graphs it has already partly seen. Source agreement between generators, nearest-neighbor distance features, and a learned embedding of candidate geometry were investigated during development but are not inputs to the final model reported here. The resulting score is a calibrated estimate of matching a released GEOM reference conformer, and it is never treated as a complete judgment of physical validity on its own, which is why Sections 4.3 and 4.4 build additional calibrated structure on top of it rather than using it directly as a selection criterion.

## B Out-of-distribution AMR

Table 6 reports the average minimum RMSD (AMR, lower is better) counterpart to Table 3’s threshold recall and precision, on the same models, datasets, and returned ensembles, now including Ref-Select (*µ* = 0.3). RefSelect has the best AMR in both columns on pepconf, and the best precision AMR on QMugs, while CovSelect keeps the best recall AMR on QMugs and both best AMR values on Platinum.

**Table 6:**
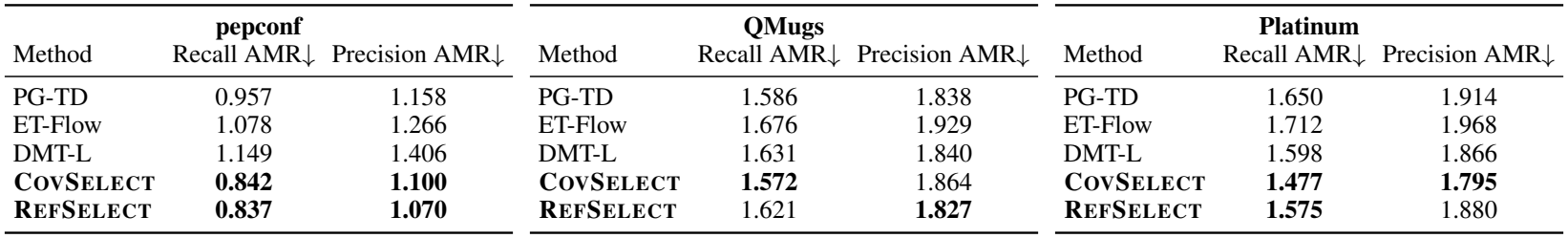
Out-of-distribution AMR (Å, lower is better), same models, datasets, and returned ensembles as Table 3. Bold marks the best value in each column.

**Table 7:** RefSelect at the 1 × and 0.5 × budgets, same *µ* grid as Table 1, on the full test set. The ‘red.’ column indicates whether redundancy correction is used.

| *μ* | red. | Budget | R mean↑ | P mean↑ | AMR-R mean↓ | AMR-P mean↓ |
|---|---|---|---|---|---|---|
| .2 | no | 1× | 85.02 | 65.00 | 0.494 | 0.663 |
| .2 | no | 0.5× | 73.53 | 70.72 | 0.630 | 0.618 |
| .3 | no | 1× | 84.33 | 67.61 | 0.498 | 0.645 |
| .3 | no | 0.5× | 72.77 | 72.05 | 0.635 | 0.608 |
| .1 | yes | 1× | 85.62 | 64.28 | 0.508 | 0.683 |
| .1 | yes | 0.5× | 73.66 | 71.13 | 0.638 | 0.625 |
| .2 | yes | 1× | 85.01 | 66.96 | 0.511 | 0.664 |
| .2 | yes | 0.5× | 73.14 | 72.46 | 0.643 | 0.617 |
| .3 | yes | 1× | 84.36 | 69.19 | 0.515 | 0.647 |
| .3 | yes | 0.5× | 72.62 | 73.64 | 0.647 | 0.609 |
| .5 | yes | 1× | 82.53 | 72.72 | 0.531 | 0.620 |
| .5 | yes | 0.5× | 71.18 | 75.70 | 0.661 | 0.594 |
| .8 | yes | 1× | 79.75 | 76.24 | 0.558 | 0.589 |
| .8 | yes | 0.5× | 68.60 | 77.93 | 0.691 | 0.575 |
| 1.2 | yes | 1× | 76.31 | 78.57 | 0.595 | 0.566 |
| 1.2 | yes | 0.5× | 65.59 | 79.95 | 0.725 | 0.556 |

## C RefSelect at Reduced Budgets

Table 7 extends the return-budget sensitivity analysis of Section 6.4 to the full RefSelect *µ* grid from Table 1, including the redundancy-corrected settings, at the 1 × and 0.5× budgets. Every setting’s precision rises and recall falls as the budget shrinks from 1 × to 0.5 ×, mirroring the pattern CovSelect shows in Table 5.

## References

Dylan M. Anstine, Roman Zubatyuk, and Olexandr Isayev. AIMNet2: a neural network potential to meet your neutral, charged, organic, and elemental-organic needs. arXiv preprint arXiv:2308.06068, 2024.

Simon Axelrod and Rafael Gómez-Bombarelli. GEOM, energy-annotated molecular conformations for property prediction and molecular generation. Scientific Data, 9(1):185, 2022.

Gabriele Corso, Yilun Xu, Valentin de Bortoli, Regina Barzilay, and Tommi Jaakkola. Particle guidance: non-i.i.d. diverse sampling with diffusion models. arXiv preprint arXiv:2310.13102, 2023.

Stephen E Farr, Stefan Doerr, Antonio Mirarchi, Francesc Sabanes Zariquiey, and Gianni De Fabritiis. Aceff: A state-of-the-art machine learning potential for small molecules. arXiv preprint arXiv:2601.00581, 2026.

Nils-Ole Friedrich, Agnes Meyder, Christina de Bruyn Kops, Kai Sommer, Florian Flachsenberg, Matthias Rarey, and Johannes Kirchmair. High-quality dataset of protein-bound ligand conformations and its application to benchmarking conformer ensemble generators. Journal of Chemical Information and Modeling, 57(3):529–539, 2017.

Teofilo F Gonzalez. Clustering to minimize the maximum intercluster distance. Theoretical computer science, 38:293–306, 1985.

Majdi Hassan, Nikhil Shenoy, Jungyoon Lee, Hannes Stark, Stephan Thaler, and Dominique Beaini. ET-Flow: Equivariant flow-matching for molecular conformer generation. In Advances in Neural Information Processing Systems, 2024.

Clemens Isert, Kenneth Atz, José Jiménez-Luna, and Gisbert Schneider. QMugs, quantum mechanical properties of drug-like molecules. Scientific Data, 9:273, 2022.

Bowen Jing, Gabriele Corso, Jeffrey Chang, Regina Barzilay, and Tommi Jaakkola. Torsional diffusion for molecular conformer generation. In Advances in Neural Information Processing Systems, 2022.

Leonard Kaufman and Peter J. Rousseeuw. Finding Groups in Data: An Introduction to Cluster Analysis. Wiley, 1990.

Hyoungrae Kim, Cheongyun Jang, Dharmendra K. Yadav, and Mi-hyun Kim. The comparison of automated clustering algorithms for resampling representative conformer ensembles with RMSD matrix. Journal of Cheminformatics, 9:21, 2017. doi: 10.1186/s13321-017-0208-0.

Sunghwan Kim, Evan E. Bolton, and Stephen H. Bryant. PubChem3D: conformer ensemble accuracy. Journal of Cheminformatics, 5:1, 2013. doi: 10.1186/1758-2946-5-1.

Zhiyuan Liu, Yanchen Luo, Han Huang, Enzhi Zhang, Sihang Li, Junfeng Fang, Yaorui Shi, Xiang Wang, Kenji Kawaguchi, and Tat-Seng Chua. NExT-Mol: 3d diffusion meets 1d language modeling for 3d molecule generation. In International Conference on Learning Representations, 2025.

Leon B. Lucy. An iterative technique for the rectification of observed distributions. The Astronomical Journal, 79:745–754, 1974.

George L. Nemhauser, Laurence A. Wolsey, and Marshall L. Fisher. An analysis of approximations for maximizing submodular set functions—i. Mathematical Programming, 14:265–294, 1978.

Philipp Pracht, Stefan Grimme, Christoph Bannwarth, Fabian Bohle, Sebastian Ehlert, Gereon Feldmann, Johannes Gorges, Marcel Müller, Tim Neudecker, Christoph Plett, et al. Crest—a program for the exploration of low-energy molecular chemical space. The Journal of Chemical Physics, 160(11), 2024.

Viki Kumar Prasad, Alberto Otero-de La-Roza, and Gino A DiLabio. Pepconf, a diverse data set of peptide conformational energies. Scientific data, 6(1):180310, 2019.

William Hadley Richardson. Bayesian-based iterative method of image restoration. Journal of the Optical Society of America, 62(1):55–59, 1972.

